# Cross-Domain Transfer Learning from Peptides to Lipids Using a Multi-Property Fine-Tuned LLM

**DOI:** 10.64898/2026.01.06.697904

**Authors:** Uchenna Alex Anyaegbunam, David Teschner, Thierry Schmidlin, Andreas Hildebrandt, Jo-hannes U Mayer, Maximilian Sprang, Miguel A. Andrade-Navarro

## Abstract

Accurate knowledge of liquid chromatography retention time (RT) is essential for confident compound identification in metabolomics and lipidomics. Yet, it is often constrained by the scarcity of experimental data for many molecular classes. Current workflows depend on experimental RT libraries, which are time-consuming to build and limited to previously observed compounds. Here, we present a transfer learning pipeline that leverages large, publicly available peptide datasets to enable accurate lipid RT prediction in data-sparse scenarios. We first show that a ChemBERTa language model, when pre-trained on peptides with a multi-task objective (predicting both RT and fundamental RDKit molecular descriptors), learns a more robust and generalizable chemical representation than a single-task (RT-only) model. This multi-property pre-training yielded superior generalization in lipids, achieving test R^2^ values of 0.842 against 0.814 (RT-only).

Crucially, transferring this peptide-based model to lipid data provided a pronounced advantage in data-sparse scenarios. When fine-tuned on only 5% of available lipid data, the transferred model improved the median test R^2^ by +0.234 over a model trained from scratch. Significant benefits persisted at intermediate data scales (50–75%), with performance converging only when 100% of the lipid data was used. Notably, the pre-trained model never underperformed the baseline, exhibiting more stable training across all data scales.

These results demonstrate that multi-property pre-training guides language models towards chemically meaningful representations that support better RT prediction in different molecular domains. Furthermore, peptide-based pre-training facilitates cross-domain transfer of chemical properties to lipid. Our work provides a practical, scalable strategy to mitigate data scarcity in lipidomics by transferring knowledge from data-rich peptide databases, offering a computational alternative to extensive experimental library generation and enabling more confident identification in small-scale omics studies.

## 1. Introduction

Accurately predicting molecular properties from chemical structure is fundamental to computational bioinformatics and drives advancements in fields such as drug discovery [1], metabolomics [2], and systems biology. Among these properties, liquid chromatography retention time (RT) is a major analytical metric. RT is essential for separating, identifying, and characterizing complex mixtures of compounds, such as metabolites and lipids, in biological samples [3,4]. Computational prediction of RT can dramatically accelerate the annotation of unknowns in untargeted omics studies, which remain a major bottleneck [5]. This challenge is particularly observed in lipidomics and metabolomics, where chemical diversity is high, but experimentally measured RTs are available for only a small fraction of known molecules. However, a persistent limitation is the scarcity of large, high-quality, labeled experimental datasets for many molecular classes [6,7].

Transfer learning (TL) has emerged as a powerful strategy to overcome data scarcity [8,9]. Knowledge gained from a data-rich source task (for example, in bottom-up proteomics, predicting peptide properties such as retention time, ion-mobility, or fragmentation patterns are well established, and rich datasets exist to train predictors for those tasks [10, 11]) is transferred to improve learning on a data-poor target task (such as predicting these properties for lipids) [12,13]. The underlying hypothesis is that neural networks can learn fundamental, domain-independent chemical principles: relationships between structure, polarity, lipophilicity, size, etc., during initial training, and these learned representations can be transferred across related biochemical domains [14,15]. Recent work in proteomics has provided strong empirical support for this hypothesis. For example, the DeepLC framework demonstrated that peptide RT prediction models pretrained on large datasets can be efficiently adapted via transfer learning to new chromatographic conditions and even to peptides bearing chemically distinct modifications, without retraining from scratch [16]. These results established transfer learning as a robust mechanism for adapting chromatographic predictors under distributional shift. However, systematic investigation of cross-domain transfer, particularly from peptides to chemically distinct molecular classes such as lipids, remains limited.

At the same time, large language models (LLMs) have revolutionized representation learning in chemistry. Models like ChemBERTa are pre-trained on a vast amount of SMILES strings, treating chemical structures as a language [17,18]. Self-supervised pre-training allows the model to learn rich, contextual embeddings of atoms and functional groups, effectively capturing the “syntax and semantics” of chemistry without labeled data [19,20]. Such models serve as versatile “foundation models” that can be fine-tuned to specific downstream tasks with relatively little data [21,22]. In metabolomics, transformer-based architectures pretrained on large molecular corpora have already demonstrated strong performance in RT prediction and transfer across chromatographic systems [23, 24].

Simply fine-tuning a base chemical LLM on a single target task (e.g., RT prediction) can be effective, but in extremely data-sparse scenarios it remains prone to overfitting and may fail to learn the most robust, generalizable features [25]. Multi-task learning (MTL) provides a compelling solution by training one model on multiple related tasks simultaneously [26,27]. In molecular property prediction, augmenting the primary task (RT) with auxiliary tasks (such as predicting fundamental physicochemical descriptors) imposes a strong inductive bias [28, 29]. The model’s internal representation, in turn, then encodes a better understanding of chemistry. For example, jointly predicting RDKit descriptors [30] like molecular weight, polar surface area, hydrogen-bond donors/acceptors, rotatable bonds, and aromatic ring count pushes the model towards learning structural principles that govern both chromatographic behavior and core molecular properties. The resulting learned features tend to be more robust and transferable than those learned from a single task alone. This strategy aligns with recent trends in metabolomics RT prediction, where combining structural representations with auxiliary molecular information has been shown to improve transferability across experimental conditions [23].

Notably, peptides and lipids, despite differences in polymeric versus modular structures, are governed by shared physicochemical principles (e.g., hydrophobicity, charge, volume) that dictate their separation in reversed-phase liquid chromatography [31,32]. A model that learns to predict peptide RT alongside its fundamental descriptors may therefore develop feature representations that are directly relevant to lipids. This strategy aligns with the vision of building “foundation models for science” that bridge data-rich and data-poor domains [33,34]. Despite this potential, there has been little systematic investigation of whether multi-property pre-training on peptide data yields features that transfer effectively to lipid RT prediction. In this study, we propose and evaluate a novel pipeline for lipid RT prediction under data-limited conditions. Our approach is founded on two main hypotheses: (1) A ChemBERTa LLM fine-tuned with a multi-task objective on peptide data (predicting both RT and seven RDKit-derived descriptors) will learn chemical representations that are superior and more transferable than those from a model fine-tuned on peptide RT alone. (2) These enriched representations will enable cross-domain transfer to lipid RT prediction, with benefits that are most pronounced in low-data domains.

To test these hypotheses, we first develop a multi-property model (ChemBERTa+RDKit) and a single-task baseline model (ChemBERTa-RT) using a large public peptide dataset. We demonstrate that the multi-property model achieves better generalization on both peptide and lipid RT prediction compared to the single-task model. We then use the multi-property peptide model as a pre-trained source for transfer learning to lipids, systematically fine-tuning it on subsets of lipid RT data ranging from 5% to 100%. We compare its performance to baseline models trained from scratch on the same lipid subsets. Our results provide clear, practical guidelines for deploying transfer learning in resource-constrained lipidomics: we observe drastic gains when lipid data is very limited (about 2% of the peptide pre-training data), substantial improvements at intermediate data scales, and convergence with the full lipid data set (about 25% of the peptide pre-training data). Overall, this multi-task transfer approach leverages large public peptide datasets to solve small-scale data challenges in omics.

## 2. Methods

### 2.1 Datasets and preprocessing

Peptide retention time data: We used a large-scale peptide RT dataset of ∼250,000 entries from the ProteomeTools project [35]. The processed data was accessed via the ProteomeTools repository on ProteomicsML [36], which provides standardized training/test splits for machine learning. Each example includes a SMILES string for a unique peptide and an experimental LC-MS/MS retention time. Lipid retention time data: For the target task, we used a lipid RT dataset derived from the METLIN SMRT database [5]. This dataset contains LC-MS retention times for diverse lipid molecules. We extracted canonical SMILES and RT values, and we constructed standardized splits (training and test) with 63,163 and 7,798 molecules, respectively. To simulate data-sparse scenarios, the lipid training set was randomly subsampled (without replacement) at fractions of 5%, 25%, 50%, 75%, and 100%. Molecular descriptors: For both peptide and lipid molecules, we computed seven fundamental physicochemical descriptors from the SMILES using the RDKit cheminformatics toolkit [30]. The descriptors were molecular weight (MW), topological polar surface area (TPSA), number of hydrogen bond donors (HBD), number of hydrogen bond acceptors (HBA), rotatable bond count, aromatic ring count, and heavy atom count. These descriptors served as auxiliary targets in the multi-task learning framework.

Data scaling: All target variables (RT and the seven descriptors) were normalized to [0, 1] using MinMaxScaler from scikit-learn [37], fit on the training data. This normalization is particularly important in chromatography, as absolute RT values strongly depend on experimental conditions such as gradient length, which can range from a few minutes to over two hours across studies. In the transfer learning experiments, the lipid RT values were scaled using the parameters learned from the peptide RT training data. This alignment ensured that the lipid RT distribution matched the pre-trained model’s output scale, which is critical for effective transfer.

### 2.2 Model architecture and pre-training

We based our models on the ChemBERTa-zinc-base-v1 transformer encoder [17], which is a variant of RoBERTa [38] pretrained on the ZINC15 database [39]. This model has 12 transformer layers, a hidden embedding size of 768, and ∼86M parameters. The special CLS token’s output from the final layer serves as the pooled molecular embedding. All ChemBERTa models used were fine-tuned starting with the default weights.

We implemented two model variants:

1. Single-Task Baseline (ChemBERTa-RT): The pretrained ChemBERTa encoder feeds into a single linear output node for RT prediction only.
2. Multi-Property (ChemBERTa+RDKit): The ChemBERTa encoder feeds into a linear layer producing an 8-dimensional output (one for RT plus one for each of the seven RDKit descriptors). This model is trained to jointly regress RT and descriptors in a multi-task setup.

For the multi-property model, the loss is the sum of mean squared errors (MSE) across all eight outputs. The code for model definition and training is available at our public GitHub repository (github.com/uchealex/chembedding). In all cases, we used the CLS token embedding as input to the regression head, since this token’s representation aggregates information from the entire SMILES sequence via self-attention.

### 2.3 Training procedures

Multi-task pre-training on peptides: We first trained the ChemBERTa+RDKit model on the full peptide dataset. Training was run for 15 epochs, minimizing the summed MSE loss across the eight outputs. We used the AdamW optimizer [40] with a learning rate of 2.5×10^−5^, batch size of 16, and standard linear warmup/decay schedules. For comparison, we also trained a single-task ChemBERTa-RT model (same hyperparameters but only one output).

Transfer learning to lipids: We used the trained multi-property peptide model as the source for lipid RT prediction. Specifically, we created a new target model by copying the transformer encoder weights from the peptide model and randomly reinitializing a fresh regression head (single output for RT). This model was then fine-tuned on each lipid subset (5%, 25%, 50%, 75%, 100% of data). We used the same hyperparameters as above (AdamW, lr=2.5e-5, 15 epochs, batch=16, warmup). During fine-tuning, we updated all encoder weights to allow the learned peptide features to adapt to the lipid domain.

For comparison, we also trained baseline models on each lipid subset from scratch (randomly initialized ChemBERTa-zinc-base-v1, single RT output). These baselines used the same architecture and hyperparameters but no peptide pre-training, to measure performance without transfer.

### 2.4 Evaluation metrics and statistical analysis

We evaluated model performance on the held-out lipid test set using the coefficient of determination (R^2^) and Mean Absolute Error (MAE). All evaluation metrics were computed after transforming model predictions and targets from the normalized [0,1] space back to their original, raw RT scale, ensuring that reported errors reflect experimentally meaningful units. For the multi-task peptide model, we report R^2^ and MAE for each of the eight outputs. For all lipid RT models (transfer and baseline), we report R^2^ and MAE for RT only. Each model was trained for 15 epochs. At the end of each epoch, we recorded R^2^ and MAE on the test set. This yielded 15 performance values per model run. To robustly summarize performance and stability, we report the median and interquartile range (IQR) (25th–75th percentile) of these epoch-wise metrics [41]. As recommended by Bouthillier et al., reporting median and IQR provides a clear measure of central tendency and variability of the training outcomes.

All models were implemented in PyTorch [42] (v2.0) and the HuggingFace Transformers library [43] (v4.30). Training was conducted on NVIDIA Tesla GPUs (T4 or V100) via Google Colab. Data processing and analysis used pandas and scikit-learn [37].

## 3. Results and Discussion

### Performance evaluation framework and dataset characteristics

We assessed model performance using R^2^ and MAE for RT prediction. The peptide dataset was used for initial model development, and the lipid dataset was subsampled (5%, 25%, 50%, 75%, 100%) to simulate varying data availability. We evaluated each model configuration (Baseline vs. Transfer at each data level) across 15 training epochs. Reported metrics are the median and IQR of epoch-wise R^2^ and MAE, providing a robust measure of performance and stability.

#### 3.1 Multi-property training with RDKit descriptors enhances peptide and lipid model generalization

We first examined whether jointly training on RT and seven computed descriptors would improve the ChemBERTa model’s predictive power. The multi-property model (ChemBERTa+RDKit) was trained to predict RT and the seven RDKit descriptors (MW, TPSA, HBD, HBA, rotatable bonds, aromatic rings, heavy atoms computed from SMILES. We compared this to the single-task baseline trained on RT only.

On peptide RT prediction, the multi-property model showed markedly improved generalization and stability (Figure 1). It achieved a final test R^2^ of 0.757 (IQR: 0.737–0.757) versus 0.743 (IQR: 0.719– 0.767) for the baseline. The final test MAE was substantially lower (258.3 vs. 295.6, both in same normalized units). Figure 2’s epoch-wise scatter shows the multi-property model’s R^2^ values are consistently higher, and its MAE values are consistently lower and less variable. This indicates that auxiliary descriptor prediction acts as a regularizer, guiding the model to learn a more robust, generalizable representation of peptide chemistry.

**Figure 1.**
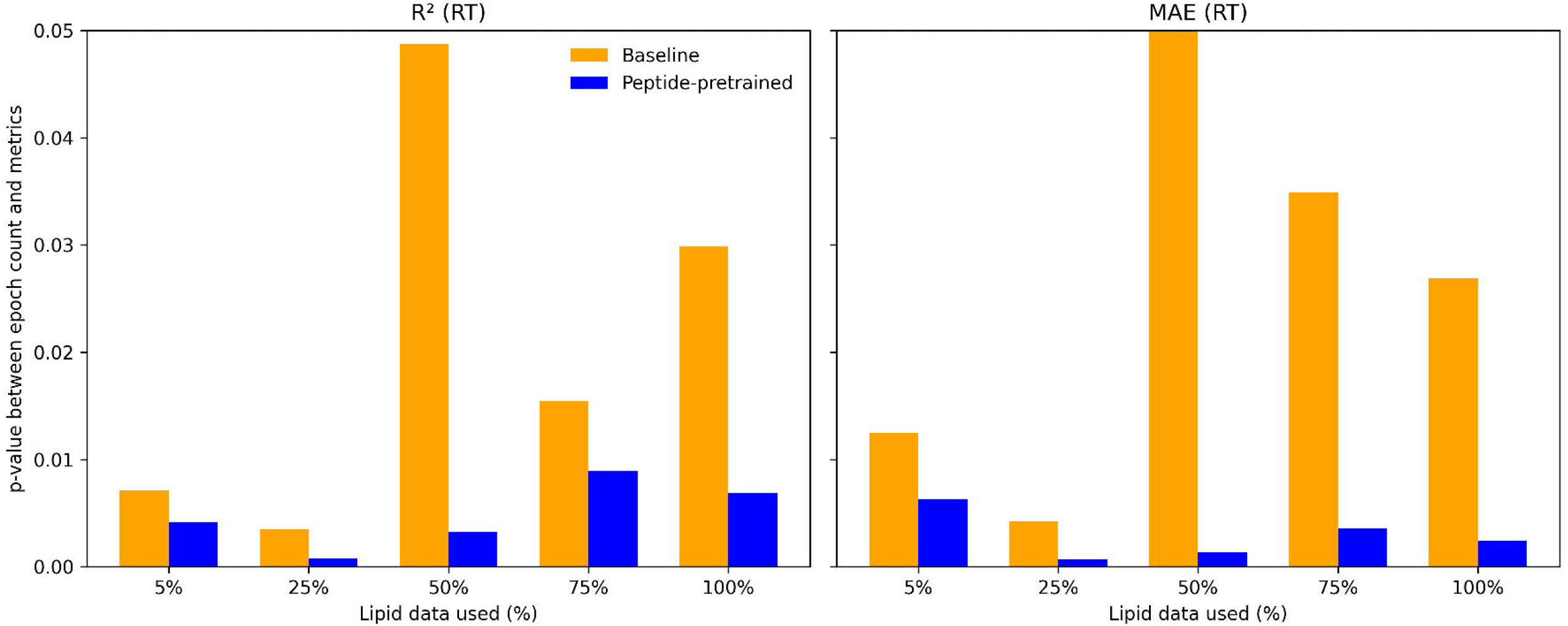
Multi-property training improves peptide retention time prediction. (A) Epoch-wise scatter plot of test versus train R^2^. The ChemBERTa+RDKit model (blue) shows a consistent upward shift in test R^2^ compared to the ChemBERTa baseline (orange), reflecting enhanced predictive performance on independent peptide data. (B) Epoch-wise scatter plot of test versus train MAE. The ChemBERTa+RDKit model (blue) achieves lower test MAE with reduced epoch-to-epoch variability, confirming that joint training with molecular descriptors yields a more reliable and generalizable model for peptide retention time.

**Figure 2.**
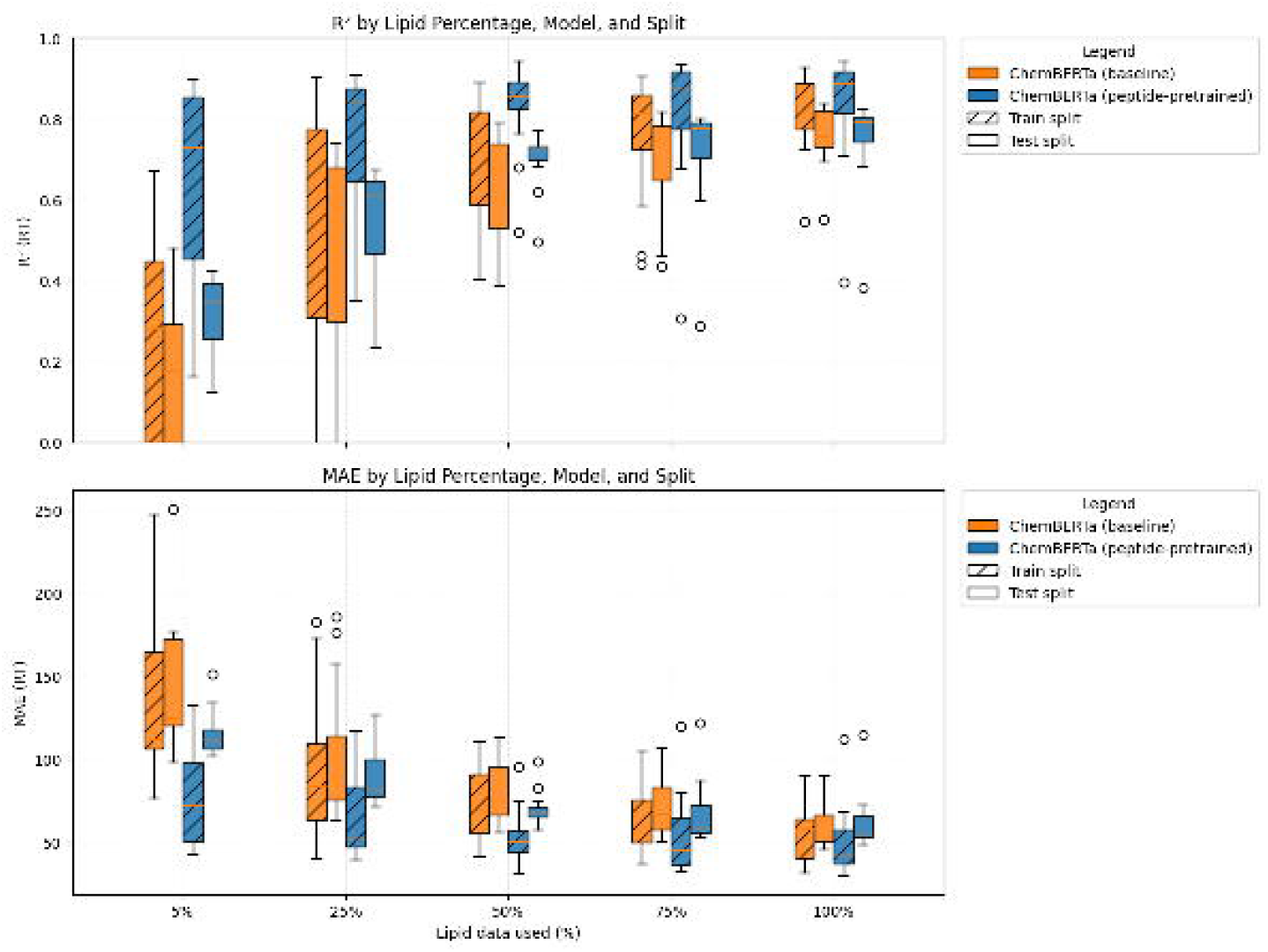
Multi-property training improves lipid retention time prediction. (A) Scatter plot of test versus train R^2^ for each of the 15 training epochs. Each point represents model performance at one epoch. The ChemBERTa+RDKit model (blue) achieves higher test R^2^ values for comparable train R^2^ values than the ChemBERTa baseline (orange), indicating better generalization. (B) Scatter plot of test versus train MAE (Mean Absolute Error, in seconds) for each epoch. Points for the ChemBERTa+RDKit model (blue) are clustered toward lower test MAE values, demonstrating more accurate and stable predictions on unseen lipid data compared to the baseline (orange). The multi-property model leverages RDKit-derived descriptors (MW, TPSA, HBD, HBA, etc.) as auxiliary tasks to learn a more robust molecular representation.

We observed a similar advantage on lipid RT prediction (Figure 2). The ChemBERTa+RDKit model attained a higher median test R^2^ (0.842, IQR: 0.817–0.847) than the baseline (0.820, IQR: 0.786– 0.819), and a lower median test MAE (45.07 vs. 52.78). Its epoch-wise results cluster at higher R^2^ and lower MAE with tighter spread. The consistent improvement across both domains strongly supports the MTL paradigm. Jointly predicting auxiliary descriptors provides a rich inductive bias, forcing the model to build a comprehensive, physically grounded molecular representation. The model thereby learns underlying structural principles rather than memorizing dataset-specific correlations. In summary, augmenting RT prediction with fundamental descriptors significantly improves model accuracy and robustness for both peptides and lipids.

#### 3.2 Transfer of a multi-property peptide model drastically improves lipid RT prediction with minimal data

Next, we tested transfer learning from peptides to lipids using the multi-property model as the source. The encoder weights from the ChemBERTa+RDKit peptide model were transferred to initialize a lipid RT model, which was then fine-tuned on subsets of lipid data. We compared this to baseline models trained from scratch on the same subsets.

Under extreme data scarcity (5% of lipids), transfer learning gave a dramatic performance boost (Figure 3, Figure 4). With only ∼3,158 lipids, the transferred model achieved a median test R^2^ of 0.393 (IQR: 0.346–0.411) versus 0.159 (IQR: –0.138–0.371) for the baseline. The median test MAE dropped from 124.3 to 105.8. Crucially, the transferred model’s performance was much more stable (narrower IQRs) across epochs. These results show that the rich, general-purpose features learned by the multi-property peptide model transfer effectively from peptides to lipids, accelerating learning when target labels are extremely scarce.

**Figure 3.**
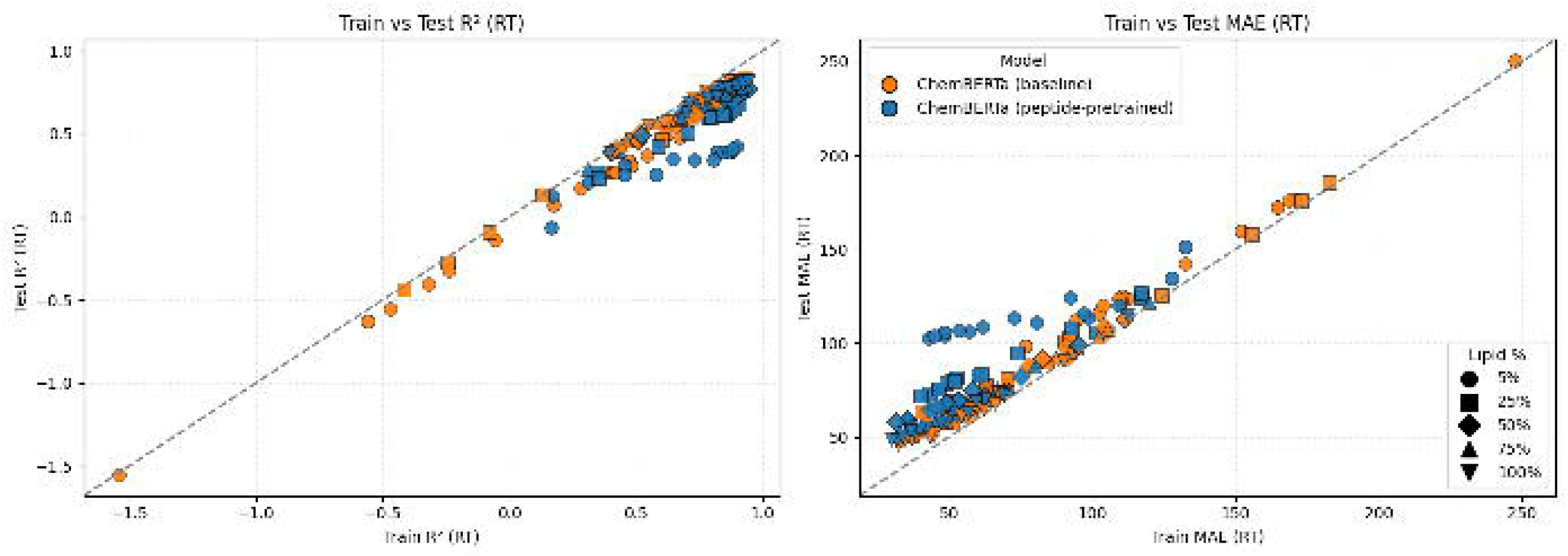
Transfer learning performance across epochs for varying lipid data amounts. (A) Test R^2^ across 15 training epochs for models trained on 5%, 25%, 50%, 75%, and 100% of the lipid dataset. The peptide-pretrained (Transfer) model (blue) consistently outperforms or matches the model trained from scratch (orange), especially in data-sparse scenarios (5%, 25%). (B) Test MAE across 15 training epochs for the same conditions. The Transfer model converges to lower error levels more quickly and maintains lower MAE, particularly when training data is scarce, highlighting the efficiency gained from pre-trained representations.

**Figure 4.**
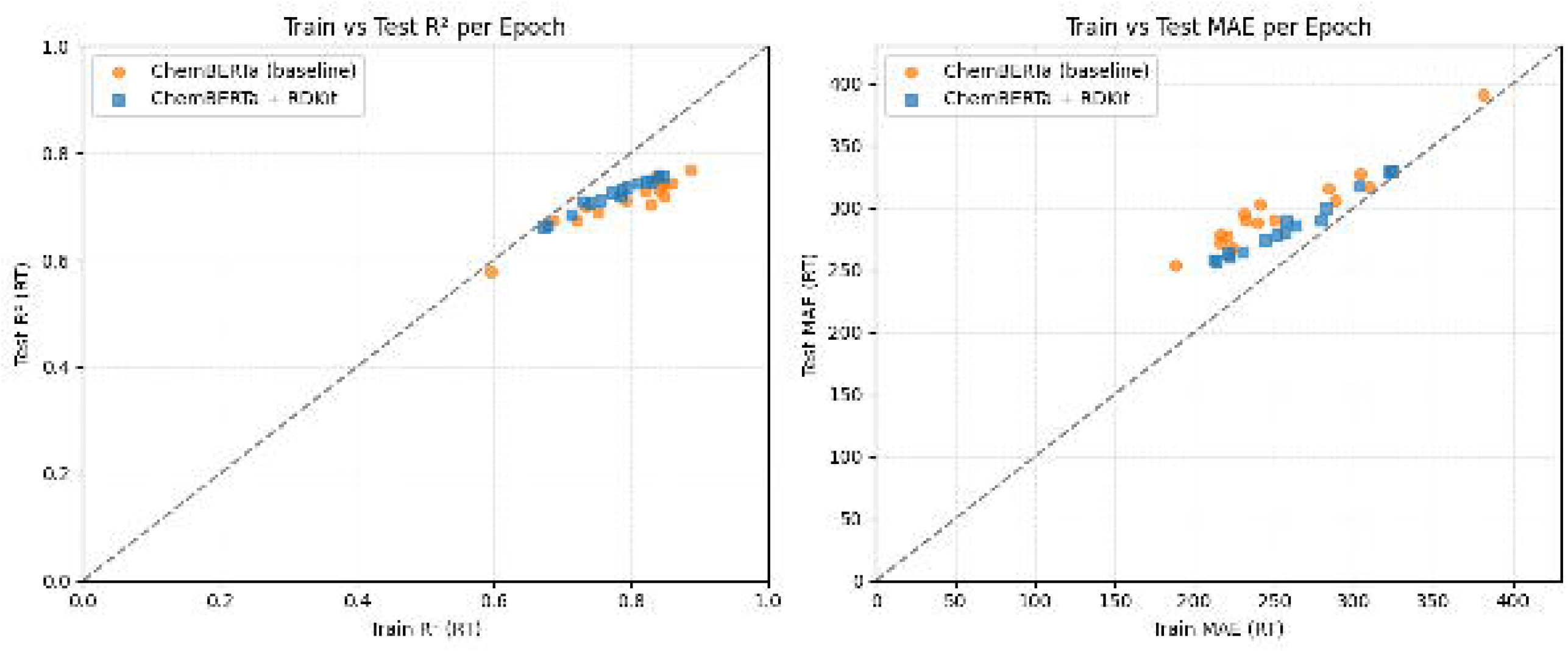
Transfer learning efficacy depends on target dataset size. (A) Boxplots (median and IQR) of test R^2^ across epochs, grouped by the percentage of lipid data used. The peptide-pretrained model (Transfer, blue) shows superior median performance and reduced variability (tighter IQR) compared to the baseline (Baseline, orange) at 5%, 25%, 50%, and 75% data. Performance converges at 100% data. (B) Boxplots of test MAE across epochs. The Transfer model achieves lower median error and greater stability (smaller IQR) than the Baseline under low-data conditions. The diminishing advantage with 100% data illustrates the saturation point of transfer benefits. Shading indicates train/test split.

#### 3.3 Transfer learning yields superior models at intermediate scales with diminishing returns at full data

At intermediate data scales (50% and 75% of lipids), the pre-trained model continued to outperform the baseline (Figure 4). With 50% of lipids, the transferred model reached a final test R^2^ of 0.771 (IQR: 0.726–0.770) versus 0.757 (IQR: 0.693–0.757) for the baseline, along with lower and less variable MAE. At 75%, transfer gave R^2^=0.797 vs. 0.762 (baseline). This demonstrates that the peptide-pretrained initialization guides optimization to a better, more stable solution even when a moderate amount of lipid data is available.

When we used 100% of the lipid training data, the performance gap narrowed. The transferred model’s final R^2^ was 0.821 (IQR: 0.798–0.824) and baseline R^2^ was 0.828 (IQR: 0.808–0.837), with similar MAEs. This convergence at full data suggests a saturation effect: with abundant lipid data, models trained from scratch can catch up. Nonetheless, across all data scales, the transferred model consistently showed reduced variability across epochs (tighter IQRs). Even when absolute gains diminish, transfer provides more reliable and reproducible training outcomes.

In summary, transfer learning from a multi-property peptide model yields substantial benefits in low- and mid-data scenarios and makes training more robust (Figure 5). This robustness is reflected in the consistently lower p-values linking epoch count to both test R^2^ and MAE for the peptide-pretrained model, indicating a stable and statistically significant improvement in performance as training progresses. In contrast, the baseline model exhibits weaker and more variable associations, particularly for MAE at intermediate data fractions, consistent with slower convergence and reduced training stability. Together, these results show that peptide-based pretraining not only improves final predictive accuracy but also accelerates learning dynamics and enhances training reliability under datalimited conditions.

**Figure 5.**
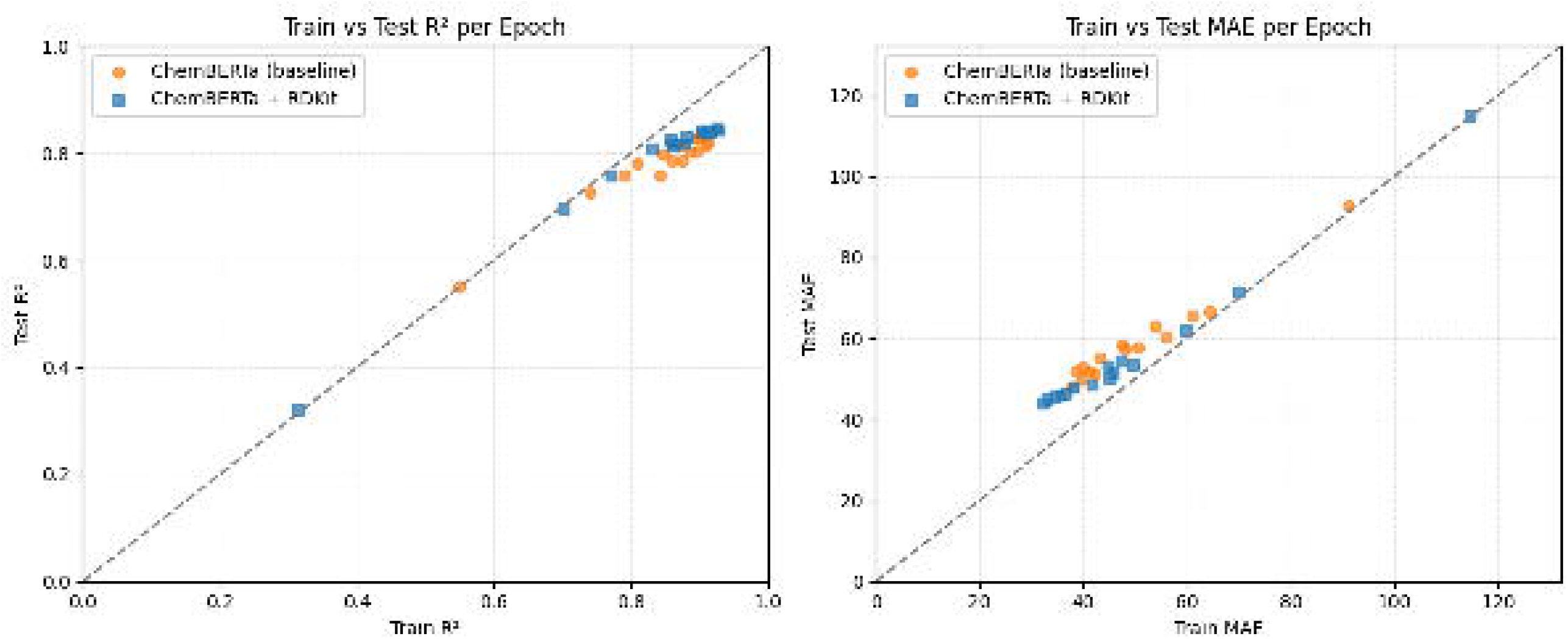
Peptide pretraining accelerates and stabilizes learning dynamics across different data-size scenarios. (A) Bar plots of Pearson correlation p-values between epoch count and test R^2^, grouped by the percentage of lipid data used. The peptide-pretrained model (Peptide-pretrained, blue) exhibits consistently lower p-values across all data fractions, indicating a strong and statistically significant association between training progression and performance improvement. In contrast, the baseline model (Baseline, orange) shows higher and more variable p-values, particularly at intermediate data sizes, reflecting slower and less stable convergence. (B) Bar plots of Pearson correlation p-values between epoch count and test MAE. The peptide-pretrained model maintains uniformly low p-values across all data-size scenarios, demonstrating robust and monotonic error reduction with increasing epochs. The baseline model displays weaker statistical associations at mid-data scenarios, consistent with reduced training stability.

## 4. Conclusion

We have demonstrated a practical pipeline for small-scale lipidomics prediction: (1) Train a multiproperty ChemBERTa model on a large peptide dataset, learning general chemical representations grounded in fundamental descriptors. (2) Transfer this model to a lipid RT prediction task under data scarcity. This approach significantly outperforms training from scratch, particularly in data-sparse scenarios. At the outset, we observe that the peptide-pretrained model is consistently more robust across all data regimes. It achieves higher accuracy at low (5%) and intermediate (50–75%) lipid data scales, while reaching performance parity with the baseline at full data availability. Importantly, no degradation in performance was observed at larger data scales, indicating an absence of negative transfer from peptide pretraining in this setting. In addition, correlation-based p-value analyses revealed that the peptide-pretrained model exhibits a more consistent and statistically significant coupling between training epochs and both test R^2^ and MAE across different data-size scenarios, indicating faster convergence and more robust training dynamics compared with the baseline model.

Our findings extend and complement prior work on transfer learning for chromatographic prediction. In proteomics, DeepLC demonstrated that pretrained peptide RT models can be efficiently adapted to new chromatographic setups and chemically distinct peptide modifications [16]. In metabolomics, models such as RT-Transformer and hybrid Transformer–LSTM architectures have shown that pretraining on large small-molecule RT datasets improves generalization across chromatographic systems [23,24]. However, these approaches still remain within a single molecular domain. Our work demonstrates effective cross-domain transfer from peptides to lipids, enabled by multi-property representation learning.

his contrasts with single-task RT models, which may overfit to dataset-specific correlations. The reduced variability and increased stability observed across all lipid data scales further indicate that multi-property pretraining produces more reliable and reproducible predictors. Even when absolute performance gains diminish at high data availability, transfer learning consistently improves training stability, as evidenced by stable loss behavior, indicating a model that is reliable and capable of maintaining predictive accuracy under varying conditions [44]. From a lipidomics and metabolomics perspective, these results are particularly relevant. RT prediction is routinely used to filter candidate annotations and to increase confidence in compound identification in untargeted workflows. Accurate RT prediction under limited data availability could reduce reliance on chemical standards and repeated experimental measurements, thereby accelerating biological interpretation. The success of peptide-to-lipid transfer further suggests that performance in chromatographic data sets is related to physicochemical principles [45], and that it can be learned from data-rich domains and reused in data-poor ones.

More broadly, our work supports the emerging paradigm of foundation models for analytical chemistry, where large, heterogeneous omics datasets are leveraged to build reusable representations that generalize across molecular classes and experimental setups. While this study focused on lipid RT prediction, the proposed strategy is likely extensible to other metabolomic properties, chromatographic modes, and molecular classes. Future work may explore bidirectional transfer between metabolites and lipids, incorporation of additional auxiliary tasks, or joint modeling across ionization modes and chromatographic conditions [46, 47].

In conclusion, this study establishes multi-task transfer learning from peptides as a robust and scalable solution for lipid RT prediction under data scarcity. By leveraging large public peptide datasets, our approach addresses a central limitation in lipidomics and metabolomics and provides a generalizable framework for chromatographic property prediction in modern omics research.

## Supplementary Data Descriptions

Supplementary Data 1: Peptide_ChemBERTa_RDkit_Boxplot_Data_Long.csv

This file contains the epoch-wise performance data used to generate Figure 1. Each row corresponds to the performance of a specific model at a given training epoch.

Columns: model (ChemBERTa (baseline) or ChemBERTa + RDKit (descriptor-augmented)), split (Train or Test), epoch (1-15), metric (R2 or MAE), value (the performance value).

Supplementary Data 2: Lipid_ChemBERTa_RDkit_Boxplot_Data_Long.csv

This file contains the epoch-wise performance data used to generate Figure 2. Each row corresponds to the performance of a specific model at a given training epoch.

Columns: model (ChemBERTa or ChemBERTa+RDKit), split (Train or Test), epoch (1-15), metric (R2 or MAE), value (the performance value).

Supplementary Data 3: TransferLearning_Lipid_Boxplot_Data_Long.csv

This file contains the aggregated epoch-wise performance data used to generate Figures 3, 4, and 5. It records the performance for both Baseline and Transfer models across all lipid data percentages. Columns: percentage (5, 25, 50, 75, 100), model (Baseline or Transfer), split (Train or Test), epoch (1-15), metric (R2 or MAE), value (the performance value).

## References

1. Lavecchia A. Machine-learning approaches in drug discovery: methods and applications. Drug Discov Today. 2015;20(3):318–331.

2. Wishart DS. Metabolomics for investigating physiological and pathophysiological processes. Physiol Rev. 2019;99(4):1819–1875.

3. Stanstrup J, Neumann S, Vrhovšek U. PredRet: prediction of retention time by direct mapping between multiple chromatographic systems. Anal Chem. 2015;87(18):9421–9428.

4. Bros K, Fabian B, Guba W, et al. Molecular representation learning for transcriptomics-guided drug discovery. Digit Discov. 2023;2:1553–1566.

5. Domingo-Almenara X, Guijas C, Billings E, et al. The METLIN small molecule dataset for machine learning-based retention time prediction. Nat Commun. 2019;10(1):5811.

6. Ching T, Himmelstein DS, Beaulieu-Jones BK, et al. Opportunities and obstacles for deep learning in biology and medicine. J R Soc Interface. 2018;15(141):20170387.

7. Yang K, Swanson K, Jin W, et al. Analyzing learned molecular representations for property prediction. J Chem Inf Model. 2019;59(8):3370–3388.

8. Pan SJ, Yang Q. A survey on transfer learning. IEEE Trans Knowl Data Eng. 2010;22(10):1345– 1359.

9. Weiss K, Khoshgoftaar TM, Wang D. A survey of transfer learning. J Big Data. 2016;3:9.

10. Buur, L. M., Declercq, A., Strobl, M., Bouwmeester, R., Degroeve, S., Martens, L., Dorfer, V., & Gabriels, R. (2024). MS2Rescore 3.0 Is a Modular, Flexible, and User-Friendly Platform to Boost Peptide Identifications, as Showcased with MS Amanda 3.0. Journal of proteome research, 23(8), 3200–3207. 10.1021/acs.jproteome.3c00785

11. Zeng, W. F., Zhou, X. X., Willems, S., Ammar, C., Wahle, M., Bludau, I., Voytik, E., Strauss, M. T., & Mann, M. (2022). AlphaPeptDeep: a modular deep learning framework to predict peptide properties for proteomics. Nature communications, 13(1), 7238. 10.1038/s41467-022-34904-3.

12. Yosinski J, Clune J, Bengio Y, et al. How transferable are features in deep neural networks? Adv Neural Inf Process Syst. 2014;27.

13. Zhuang F, Qi Z, Duan K, et al. A comprehensive survey on transfer learning. Proc IEEE. 2020;109(1):43–76.

14. Ramsundar B, Eastman P, Walters P, et al. Deep learning for the life sciences. O’Reilly Media; 2019.

15. Xu Y, Ma J, Liaw A, et al. Demystifying multitask deep neural networks for quantitative structure–activity relationships. J Chem Inf Model. 2017;57(10):2490–2504.

16. Bouwmeester, R., Nameni, A., Declercq, A., Devreese, R., Velghe, K., Gorshkov, V., Penanes, P. A., Kjeldsen, F., Rompais, M., Carapito, C., Gabriels, R., & Martens, L. (2025). DeepLC introduces transfer learning for accurate LC retention time prediction and adaptation to substantially different modifications and setups. bioRxiv. 10.1101/2025.06.01.657225

17. Chithrananda S, Grand G, Ramsundar B. ChemBERTa: large-scale self-supervised pretraining for molecular property prediction. arXiv. 2020;2010.09885.

18. Wang S, Guo Y, Wang Y, et al. SMILES-BERT: large scale unsupervised pre-training for molecular property prediction. Proc 10th ACM Int Conf Bioinform Comput Biol Health Inform. 2019:429–436.

19. Devlin J, Chang M-W, Lee K, Toutanova K. BERT: pre-training of deep bidirectional transformers for language understanding. Proc NAACL-HLT. 2019:4171–4186.

20. Rives A, Meier J, Sercu T, et al. Biological structure and function emerge from scaling unsupervised learning to 250 million protein sequences. Proc Natl Acad Sci USA. 2021;118(15):e2016239118.

21. Bommasani R, Hudson DA, Adeli E, et al. On the opportunities and risks of foundation models. arXiv. 2021;2108.07258.

22. Zvyagin M, Brace A, Hippe K, et al. GenSLMs: genome-scale language models reveal SARS-CoV-2 evolutionary dynamics. bioRxiv. 2022.

23. Xue, J., Wang, B., Ji, H., & Li, W. (2024). RT-Transformer: retention time prediction for metabolite annotation to assist in metabolite identification. Bioinformatics (Oxford, England), 40(3), btae084. 10.1093/bioinformatics/btae084

24. Mazraedoost, S., Sedigh Malekroodi, H., Žuvela, P., Yi, M., & Liu, J. J. (2025). Prediction of Chromatographic Retention Time of a Small Molecule from SMILES Representation Using a Hybrid Transformer-LSTM Model. Journal of chemical information and modeling, 65(7), 3343–3356. 10.1021/acs.jcim.5c00167

25. Chen L, Tan X, Wang D, et al. Multitask learning with graph neural networks for molecular property prediction. ACS Omega. 2021;6(46):30815–30823.

26. Caruana R. Multitask learning. Machine Learning. 1997;28(1):41–75.

27. Zhang Y, Yang Q. A survey on multi-task learning. IEEE Trans Knowl Data Eng. 2021;34(12):5586–5609.

28. Gan, H., Fu, L., Zhou, R., Gan, W., Wang, F., Wu, X., Yang, Z., & Huang, Z. (2024). WAL-Net: Weakly supervised auxiliary task learning network for carotid plaques classification. Engineering Applications of Artificial Intelligence, 137(Part A), Article 109144. 10.1016/j.en-gappai.2024.109144.

29. Wang, Y., Xiong, J., Xiao, F., Zhang, W., Cheng, K., Rao, J., Niu, B., Tong, X., Qu, N., Zhang, R., Wang, D., Li, X., & Zheng, M. (2023). LogD7.4 prediction enhanced by transferring knowledge from chromatographic retention time, microscopic pKa and logP. Journal of cheminformatics, 15(1), 76. 10.1186/s13321-023-00754-4.

30. Landrum G. RDKit: open-source cheminformatics software. http://www.rdkit.org. 2020.

31. Kyle JE. Lipidomics in translational research and the clinical significance of lipid-based biomarkers. Transl Res. 2017;189:13–29.

32. Cajka T, Fiehn O. Toward merging untargeted and targeted methods in mass spectrometry-based metabolomics and lipidomics. Anal Chem. 2016;88(1):524–545.

33. Greener JG, Kandathil SM, Moffat L, et al. A guide to machine learning for biologists. Nat Rev Mol Cell Biol. 2022;23(1):40–55.

34. Xu Z, Wang X, Wu Z, et al. Molecular contrastive learning with chemical element knowledge graph. Proc AAAI Conf Artif Intell. 2022;36(4):3968–3976.

35. Zolg, D. P., Wilhelm, M., Schnatbaum, K., Zerweck, J., Knaute, T., Delanghe, B., Bailey, D. J., Gessulat, S., Ehrlich, H. C., Weininger, M., Yu, P., Schlegl, J., Kramer, K., Schmidt, T., Kusebauch, U., Deutsch, E. W., Aebersold, R., Moritz, R. L., Wenschuh, H., Moehring, T., … Kuster, B. (2017). Building ProteomeTools based on a complete synthetic human proteome. Nature methods, 14(3), 259–262. 10.1038/nmeth.4153.

36. Rehfeldt, T. G., Gabriels, R., Bouwmeester, R., Gessulat, S., Neely, B. A., Palmblad, M., Perez-Riverol, Y., Schmidt, T., Vizcaíno, J. A., & Deutsch, E. W. (2023). ProteomicsML: An Online Platform for Community-Curated Data sets and Tutorials for Machine Learning in Proteomics. Journal of proteome research, 22(2), 632–636. 10.1021/acs.jproteome.2c00629

37. Pedregosa F, Varoquaux G, Gramfort A, et al. Scikit-learn: machine learning in Python. J Mach Learn Res. 2011;12:2825–2830.

38. Liu Y, Ott M, Goyal N, et al. RoBERTa: a robustly optimized BERT pretraining approach. arXiv. 2019;1907.11692.

39. Sterling T, Irwin JJ. ZINC 15 – ligand discovery for everyone. J Chem Inf Model. 2015;55(11):2324–2337.

40. Loshchilov I, Hutter F. Decoupled weight decay regularization. Proc Int Conf Learn Represent. 2019.

41. Bouthillier X, Delaunay P, Bronzi M, et al. Accounting for variance in machine learning benchmarks. Proc Mach Learn Sys. 2021;3:747–769.

42. Paszke A, Gross S, Massa F, et al. PyTorch: an imperative style, high-performance deep learning library. Adv Neural Inf Process Syst. 2019;32:8024–8035.

43. Wolf T, Debut L, Sanh V, et al. Transformers: state-of-the-art natural language processing. Proc EMNLP 2020: System Demonstrations. 2020:38–45.

44. Li, J., Lin, Y., Gui, Z., & Wang, P. (2025). Inception–Attention–BiLSTM Hybrid Network: A Novel Approach for Shear Wave Velocity Prediction Utilizing Well Logging. Applied Sciences, 15(5), 2345. 10.3390/app15052345

45. Bandini, E., Castellano Ontiveros, R., Kajtazi, A., Eghbali, H., & Lynen, F. (2024). Physicochemical modelling of the retention mechanism of temperature-responsive polymeric columns for HPLC through machine learning algorithms. Journal of cheminformatics, 16(1), 72. 10.1186/s13321-024-00873-6

46. Lee, I. C. H., Tumanov, S., Wong, J. W. H., Stocker, R., & Ho, J. W. K. (2023). Integrative processing of untargeted metabolomic and lipidomic data using MultiABLER. iScience, 26(6), 106881. 10.1016/j.isci.2023.106881

47. Sibilio, P., De Smaele, E., Paci, P., & Conte, F. (2025). Integrating multi-omics data: Methods and applications in human complex diseases. Biotechnology reports (Amsterdam, Netherlands), 48, e00938. 10.1016/j.btre.2025.e00938

